# Effect of Puerarin on PI3K-AKT Signaling Pathway in Osteoclast

**DOI:** 10.1101/2022.05.25.493471

**Authors:** Yiqiu Yang, Lan Li, Na Zhao, Shanshan Kuang, Yaowen Zhang, Jisheng Xie

## Abstract

**objective:** This study intends to explore the role of PI3K-AKT signaling pathway in the effect of Puerarin on the proliferation, activity, and function of osteoclasts from the perspective of antioxidation.

**Methods:** RAW264. 7 cells were divided into control group, induction group treated with 20ng/mL M-CSF and 50ng/mL RANKL, puerarin group treated with 20ng/mL M-CSF, 50ng/mL RANKL, and 50μmol/L puerarin. The staining of osteoclasts before and after puerarin intervention was measured by TRAP staining and cell count. The changes of related molecules before and after puerarin intervention in osteoclasts were detected by real-time fluorescence quantitative PCR and Western Blot, including TRAP, MMP-9, Cathepsin K, NFATc1, PTEN, Catalase, PI3K, AKT, P-AKT(ser473), FoxO1, P-FoxO1.

**Results:** TRAP staining showed that puerarin inhibited the proliferation and differentiation of RAW264.7 cells into osteoclasts. The results of qRT-PCR and WB showed that compared with the control group, the gene expression of TRAP, MMP-9, cathepsin K and NFATc1 in the induction group was up-regulated, while the gene expression of Catalase was down-regulated. PTEN gene had no significant changes before and after puerarin intervention. The expression of P-AKT (ser473) and NFATc1 protein was up-regulated, while the expression of PI3K and AKT protein had no change. Compared with the induction group, the gene expression of TRAP, MMP-9, Cathepsin K, and NFATc1 in the puerarin group decreased, the gene expression of Catalase increased, the protein expression of PI3K and AKT remained unchanged, the protein expression of P-AKT (ser473), P-FoxO1 and NFATc1 decreased, and the protein expression of FoxO1 and Catalase increased.

**Conclusion:** Puerarin may promote the transcriptional activity of FoxO1, increase the expression of catalase protein and exert its antioxidant activity by regulating the PI3K-AKT signal pathway, so as to inhibit the proliferation and differentiation of osteoclasts.

---

Due to lower estrogen levels, osteoclast-driven bone resorption is stronger than osteoblast-driven bone production in postmenopausal individuals with osteoporosis, resulting in decreased bone mass, degradation of bone tissue microarchitecture, increased bone fragility, and increased fracture risk[1]. Oxygen radicals, also known as reactive oxygen species (ROS), are oxygen-containing reactive molecules such as hydrogen peroxide, hydroxyl radicals, and peroxides [2, 3]. It has a role in the development of osteoporosis by influencing the proliferation and differentiation of osteoblasts and osteoclasts, and is regarded a major risk factor for the disease [4-6].

Puerarin is the primary active element in isoflavones isolated from Pueraria lobata, which acts as a phytoestrogen and possesses anti-inflammatory, antioxidant, anti-tumor, and osteoporosis preventive and therapeutic properties[7,8]. Animal studies have shown that Puerarin improves the bone microenvironment by regulating short-chain fatty acid levels and repairing intestinal mucosal integrity, thereby regulating intestinal flora disorders and subsequently treating osteoporosis caused by OVX[9]. In vitro experiments have shown that puerarin induces differentiation of bone marrow stromal cells (BMSCs) into osteoblasts through the ERK1/2-Runx2 signaling pathway and inhibits osteoclast differentiation by inhibiting autophagy and proliferation of osteoclast precursors[10].

Our previous study found that Puerarin may negatively regulate the PI3K-AKT signaling pathway by elevating PTEN expression, leading to an increase in catalase and thus alleviating and treating osteoporosis[11]. Since the osteoclast-led bone resorption effect is stronger than the osteoblast-led bone formation effect in postmenopausal osteoporosis. Therefore, we propose the hypothesis that Puerarin could negatively regulate the PI3K-AKT signaling pathway by increasing PTEN expression, thereby inhibiting osteoclastogenesis and thus achieving a therapeutic effect in osteoporosis. To this end, we propose to interfere with the differentiation of RAW264.7 cells into osteoclasts through puerarin and examine the effect of puerarin on the PI3K-AKT signaling pathway in osteoclasts.

## Materials and Methods

### Experimental materials

RAW264.7 cells were obtained from the Cell Bank of the Chinese Academy of Sciences (Shanghai, China). DMEM High glucose culture medium, fetal bovine serum (FBS), 1% Penicillin-Streptomycin, Trypsin-EDTA (0.25%), phenol red, were purchased from Gibco (Carlsbad, CA, U.S.A.). QuickBlock™ Primary Antibody Dilution Buffer for Western Blot, QuickBlock™ Secondary Antibody Dilution Buffer for Western Blot, QuickBlock™ Blocking Buffer for Western Blot were supplied from Beyotime Institute of Biotechnology (Shanghai, China). ECL luminescent liquid and the mouse first antibodies against β-actin, PI3K, AKT, NFATc1, FoxO1 and Catalase were purchased from Affinity (Ohio, U.S.A.). TRAP Staining Kit was provided by Wako Pure Chemical Industries (Osaka, Japan). Total RNA Extraction Kit, HifairTM III One Step RT-qPCR Probe Kit were offered from Yeasen Biotechnology (Shanghai, China). Nunc cell culture plates were supplied from Thermo Fisher Scientific (Waltham, MA, U.S.A.). BCA Protein Assay Kit and PAGE Gel Quick Preparation Kit (10%) were supplied from Epizyme Biomedical Technology (Shanghai, China). Puerarin was provided by Chengdu Alfa Biotechnology (purity≥98% by HPLC, Chengdu, China).

### Cell Culture

RAW264.7 cells were seeded onto 24-well plates with 3×103 cells per well and grown in DMEM medium supplemented with 10% FBS and 1%(v/v) penicillin-streptomycin solution, incubated at 37°C in humidified air with 5% CO_2_. After the cells had adhered to the wall for 24 hours, they were separated into four groups: control group, induction group was treated with M-CSF (20ng/mL) and RANKL(50ng/mL), and puerarin group was treated with M-CSF (20ng/mL), RANKL(50ng/mL) and puerarin (50μmol/L). Every two days, the fluid was replaced. TRAP stain was used when osteoclasts were effectively produced, and the expression level of associated molecules was determined by qRT-PCR and Western Blot.

### TRAP staining

CTCC TRAP staining, after the cells have been cultured in the 24-well cell culture plate for the indicated number of days, the medium was ruled out. Wash the cell culture plates twice with Washing Solution No. 1, then add the fixing solution and fix for 5 minutes at room temperature. Add Washing Solution No. 2, wash 2 times, then, add TRAP staining solution and incubate for 15 minutes at 37°C in an incubator freed from light. After staining, wash twice with washing solution No. 2, examine under a microscope, and take pictures. If the osteoclast anti-tartrate acid phosphatase activity is low, the staining time can be extended appropriately to achieve the desired shade of staining under the microscope.

Wako TRAP staining, after the cells have been cultured in the 24-well cell culture plate for the indicated number of days, the medium was discarded. Wash the cell culture plates twice with PBS, then add 500μl fixing solution and left to sit on ice for 10 minutes. Pour off the fixative, washed with PBS, add 100 μl prepared TRAP staining solution to each well and leave in an incubator at 37°C for 15 minutes. After the staining was completed, the cells were washed twice with PBS and then 100μl of PBS was added to each well to prevent the sample from drying out and interfering with the observation. Finally, the cell culture plate was placed on a fluorescent inverted microscope for observation and photographic recording.

### Real-time fluorescence quantitative PCR analysis

After the cells have been cultured for the indicated number of days, total RNA is extracted according to the reagent instructions, followed by reverse transcription of the RNA into cDNA and further amplification of the target gene. The conditions for reverse transcription PCR were incubation at 42°C for 60min and heating at 70°C for 5 min. The amplification conditions were pre-denaturation at 95°C for 300s, denaturation at 95°C for 10s, and annealing/extension at 60°C for 30s, for a total of 40 cycles. All the above operations were performed on ice. The primer sequences were listed in Table 1-1.

**Table 1-1.**
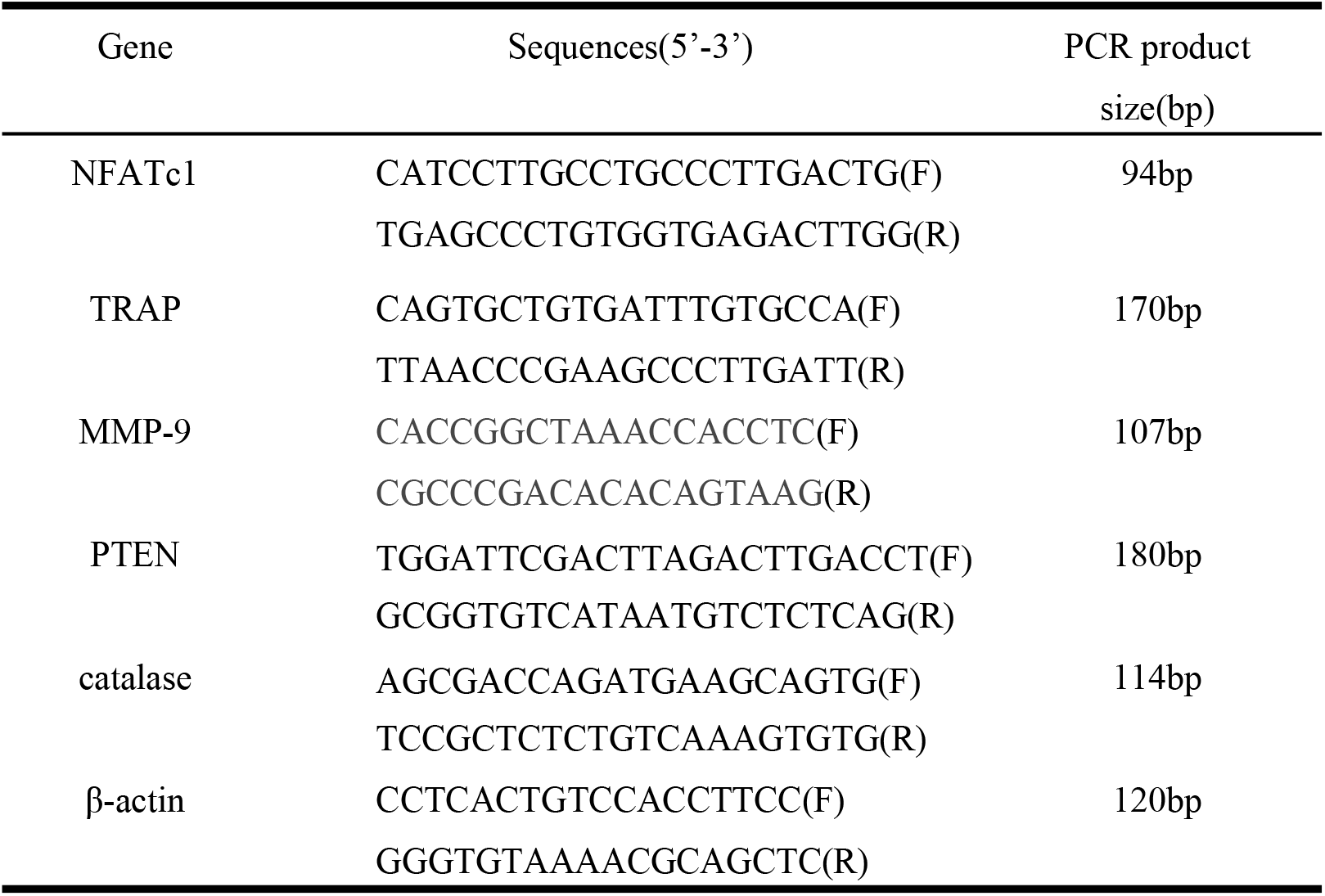
Primer sequence of target gene

### Western Blot analysis

After the cells had been cultured for the specified time, total cell proteins were collected according to the reagent instructions. The proteins were separated by 10% sodium dodecyl sulfate-polyacrylamide gel electrophoresis (SDS-PAGE) electrophoresis and the separated proteins were electrophoretically transferred to polyvinyl difluoride (PVDF)membranes. The PVDF membrane was placed in the closure solution and shaken for 25min at room temperature. Remove the PVDF membrane from the closure solution and wash it 3 times in 1×TBST for 5 min each time. Place PVDF membrane in an incubator, add primary antibody, and shake overnight at 4°C. Remove the PVDF membrane with incubated primary antibody and wash it 3 times in 1 × TBST for 5 min each time, then add the corresponding secondary antibody and incubate for 2 hours at room temperature with shaking. The PVDF membrane after incubation with secondary antibody was washed 3 times for 5 min each in 1×TBST. The prepared ECL luminescent solution was evenly dropped onto the membrane, exposed for development in a gel imaging system, and photographed for storage. Finally, the intensity of the bands was quantified by scanning densitometry by the Image J software.

### Statistical analysis

The measurement data are expressed by mean ± standard deviation(S.D.). The difference between multiple groups is compared by one-way ANOVA. When the variance is homogeneous, Fisher’s least significant difference (LSD) is used. When the variance is uneven, Dunnett’s T3 is used. *P* values < 0.05 were considered significant.

## RESULTS

### The effect of cell growth state and seed plate density of RAW264.7 on osteoclast differentiation

After inoculating RAW264.7 cells onto cell culture plates at various densities, it was discovered that their potential to develop into osteoclasts was controlled by their own growth state and the density of the seeding plate. Normal RAW264.7 cells are round, small and full with no or few tentacles. When RAW264.7 cells are hyperdifferentiated, their morphology changes, such as spindle, pentagonal, or rhomboid. At this time, RAW264.7 cells did not develop into osteoclasts even if the induction solution was increased or the time was prolonged (Fig. 1). The same is true when the cell density is too small or too large (Fig. 2).

**Fig 1.**
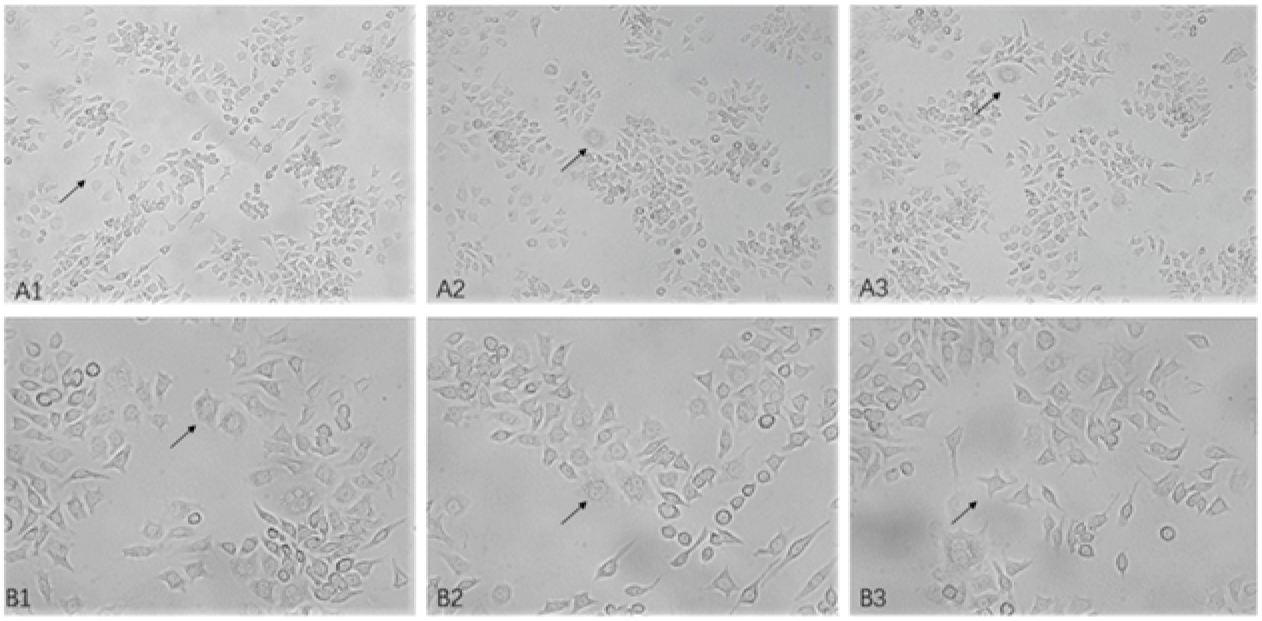
Effect of RAW264.7 cells status on osteoclast differentiation RAW264.7 cells were observed with an inverted phase contrast microscope, and the black arrows in the figure showed that the cells had differentiated into different shapes, including spindle, pentagon, and diamond. A1, A2, A3 are 10× maginification. B1, B2, B3 are 20× maginification.

**Fig.2.**
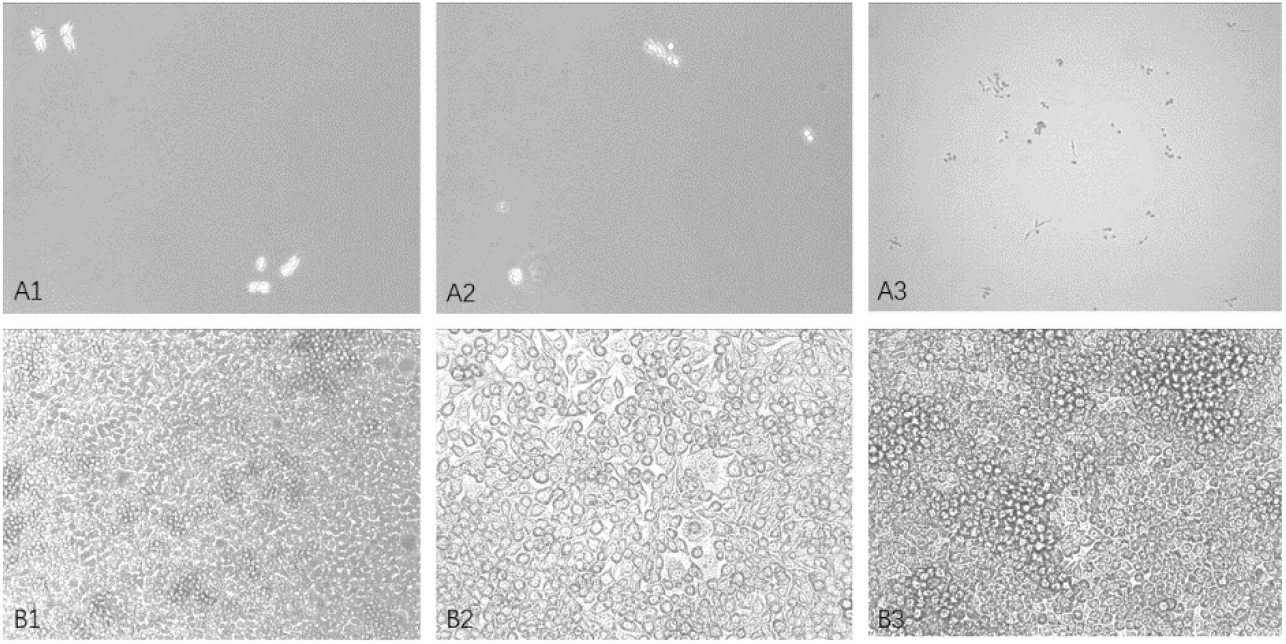
Effects of different cell densities on osteoclast differentiation RAW264.7 cells were observed by inverted phase contrast microscope, figure A1, figure A2, and figure A3 showed that the cell density was too small. figure B1, figure B2, and figure B3 showed that the cell density was too large. A1, A2, A3, B1 are 10 × maginification. B2, B3, are 20× maginification.

### Morphological observation of TRAP staining of osteoclasts

RAW264.7 cells were seeded at 3×103 cells per well in 24-well plates and cultivated for 4 days in induced differentiation medium. After TRAP staining, it can be seen that the osteoclasts were stained pink and the positive rate is higher. In mature osteoclasts, several nuclei may be observed, and the cytoplasm is plentiful (Fig. 3).

**Fig. 3.**
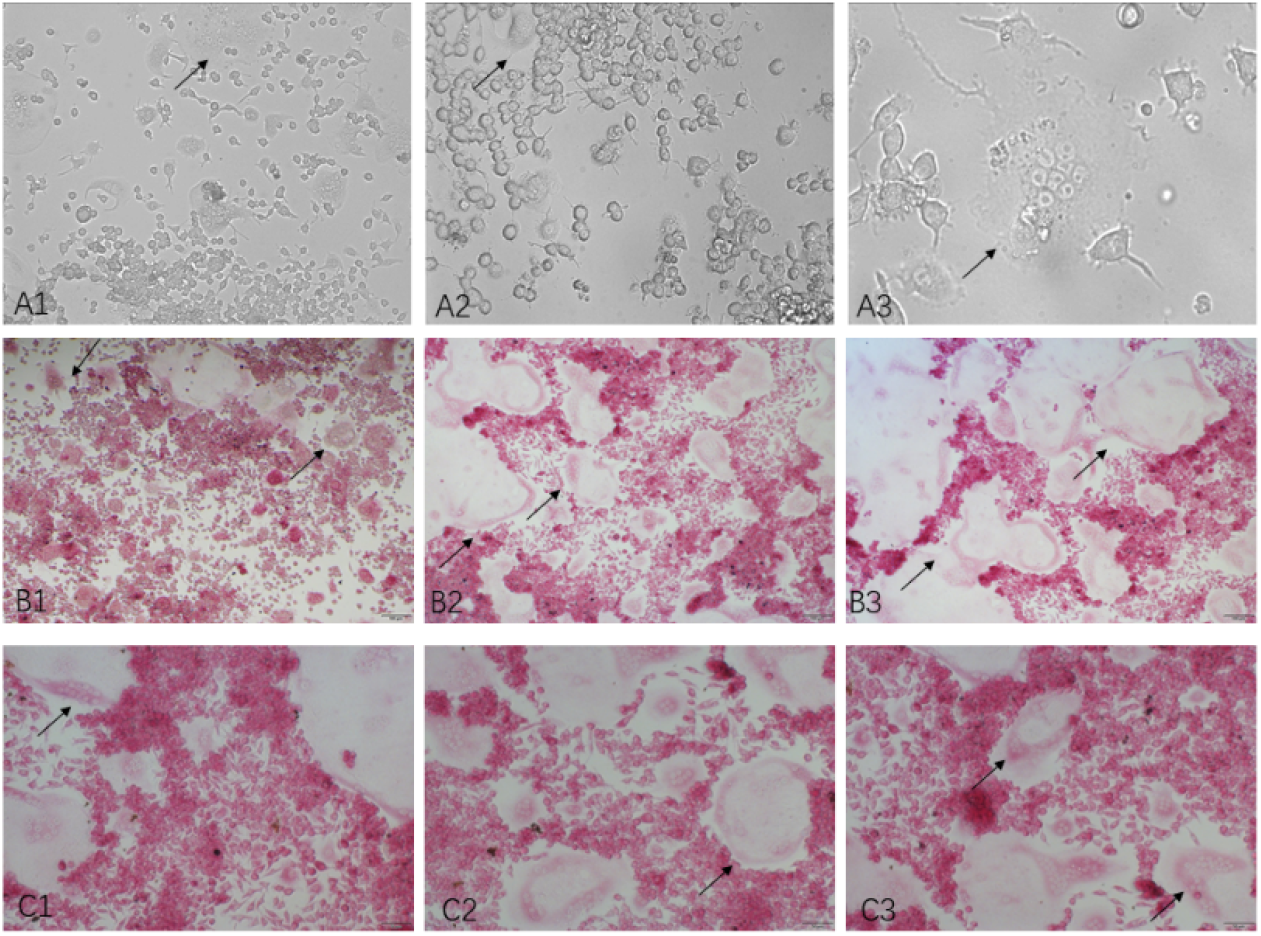
TRAP staining was used to label mature osteoclasts. Mature osteoclasts are rich in cytoplasm and possess multiple nuclei. Developed osteoclasts stained pink by Wako TRAP staining. A1, B1, B2, B3, C1, C2, C3, are 10× maginification, A2 20× is maginification, A3 is 40× maginification.

### The effect of puerarin on the differentiation of osteoclasts

In order to determine the effect of puerarin on the differentiation of osteoclasts, we seeded the cells into 24-well plates at the density of 3×103 cells per well, and divided the cells into Control group, RANKL group, and Pue group. After 4 days of culture, the cells were detected by TRAP staining. The Control group was not stained, indicating that no osteoclasts were formed in the plate, the cells in the RANKL group were stained purple, indicating that more osteoclasts were formed in the plate, while the Pue group had fewer cells stained purple, and the changes in cell morphology were less obvious than those in the RANKL group. The above results showed that puerarin inhibited the ability of RAW264.7 cells to differentiate into osteoclasts, and the difference was statistically significant (*p*<0.01) (Fig. 4).

**Fig.4.**
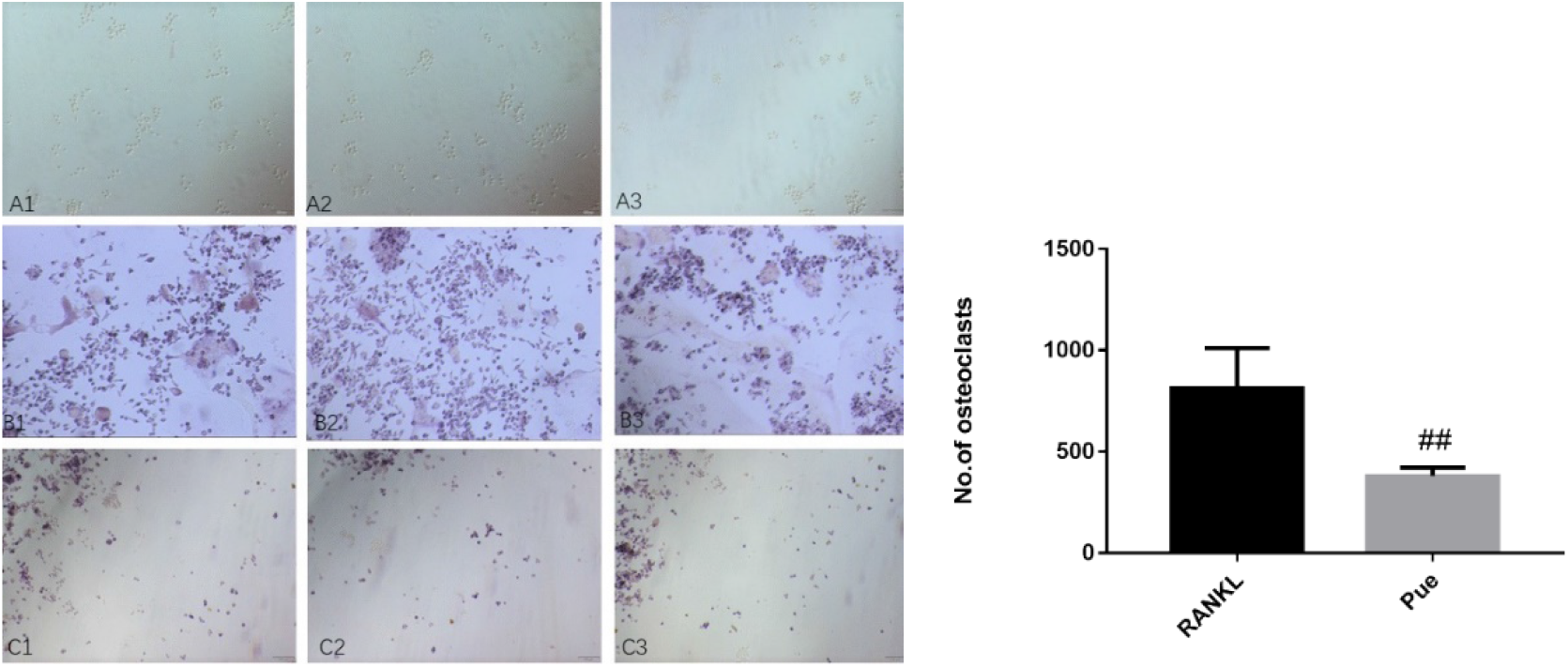
The effect of Puerarin treatment on differentiation of RAW264.7 cells into osteoclasts TRAP staining was performed on the cells and the number of TRAP+ cells was counted. Bar chart of the effect of Puerarin treatment to osteoclast number. The number of positive osteoclasts decreased significantly after puerarin intervention. ##*p*<0.001 vs induction group.

### Effects of puerarin on TRAP, MMP-9, Cathepsin K

In order to determine the effect of puerarin on the activity of osteoclasts, the mRNA expression levels of osteoclast-specific genes TRAP, MMP-9 and cathepsin K were detected by real-time fluorescence quantitative PCR. These three enzymes are specifically secreted by osteoclasts and are an important indicator reflecting the activity of osteoclasts. Compared with the Control group, the gene expressions of TRAP, MMP-9 and cathepsin K were up-regulated in the RANKL group. Compared with the RANKL group, the expression of TRAP, MMP-9 and cathepsin K was down-regulated after puerarin intervention. The above studies show that puerarin can inhibit the differentiation of RAW264.7 cells into osteoclasts (Fig. 5).

**Fig.5.**
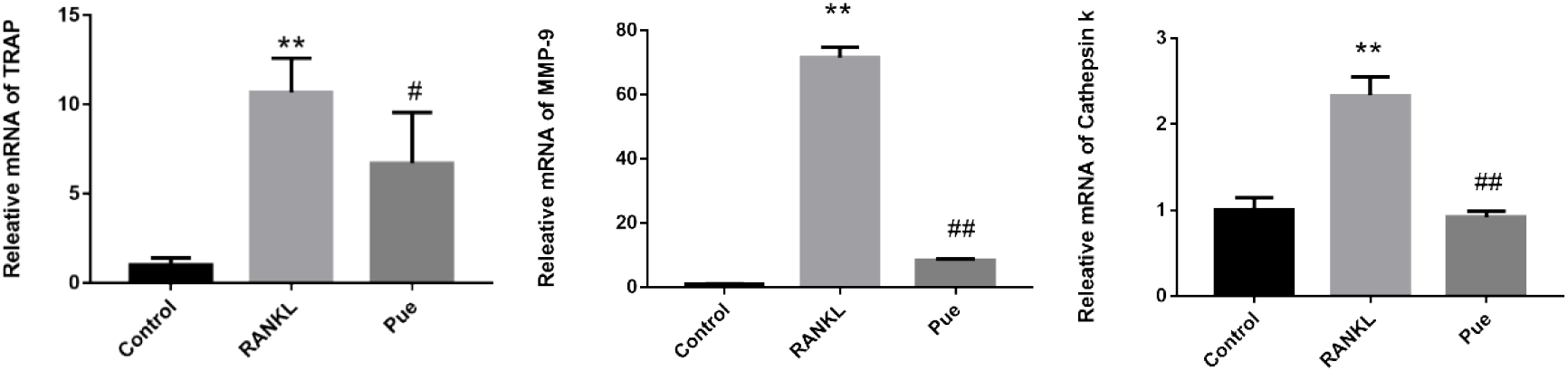
qRT-PCR measurement of Puerarin on TRAP, MMP-9, and cathepsin K. Compared with the control group, the gene expression of TRAP MMP-9 and Cathepsin K increased in the induced group. Compared with the induction group, the gene expression of TRAP, MMP-9, Cathepsin K decreased in the puerarin group.** *P* < 0.01 vs control group. # *P* < 0.05, ## *P* < 0.01 vs induction group.

### Effects of puerarin on the gene expression of PTEN, NFATc1, Catalase

We used real-time fluorescence quantitative PCR to detect the effect of puerarin on the gene expression of PTEN, NFATc1 and Catalase. Catalase gene expression was down-regulated in the RANKL group compared to the Control group, whereas NFATc1 gene expression was up-regulated. In the Pue group, compared to the RANKL group, Catalase gene expression was up-regulated and NFATc1 gene expression was down-regulated, and the difference was statistically significant (*p*<0.01). PTEN, on the other hand, did not alter appreciably before or after puerarin treatment (Fig. 6).

**Fig.6.**
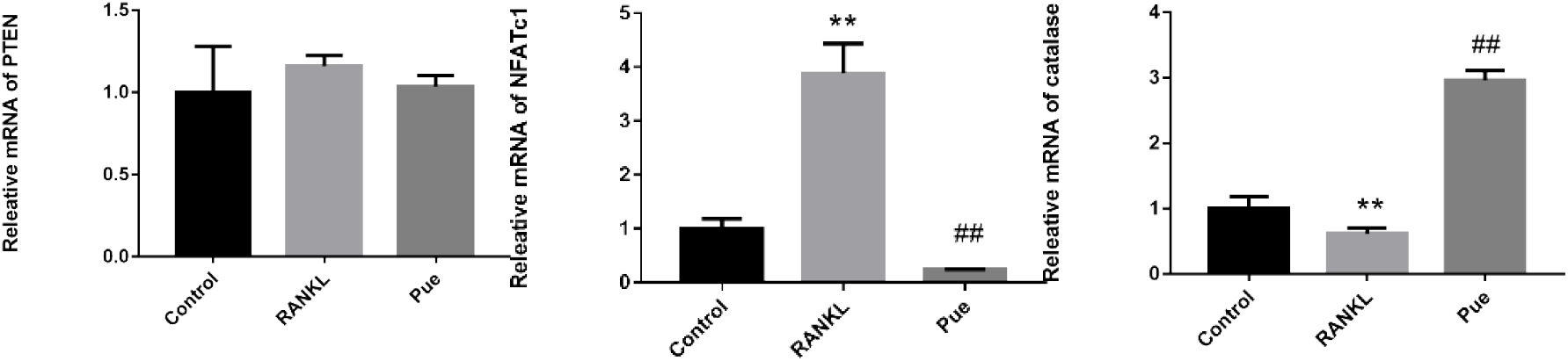
qRT-PCR measurement of Puerarin on PI3K-AKT signaling pathway Compared with the control group, the gene expression of NFATc1 in the induced group was up-regulated, while the gene expression of catalase was down-regulated. Compared with the induction group, the gene expression of NFATc1 was down-regulated in the puerarin group, while the catalase gene expression was up-regulated. The expression of PTEN gene did not change significantly before and after puerarin intervention.** *P* < 0.01 vs control group, ## *P* < 0.01 vs induction group.

### Effects of puerarin on PI3K-AKT signaling pathway

To further investigate the effect of puerarin on the PI3K-AKT signaling pathway in osteoclasts, we examined the expression of relevant proteins by Western Blot analysis. The induced osteoclasts expressed more p-AKT (ser473) and NFATc1 proteins than the control group, whereas the levels of PI3K and AKT proteins remained unchanged. The protein levels of PI3K and AKT did not alter following puerarin intervention compared to the RANKL group, however the expressions of p-AKT (ser473) and NFATc1 protein were down-regulated. This indicates that the inhibition of osteoclast differentiation by puerarin may be related to the inhibition of the activation of PI3K-AKT signaling pathway (Fig. 7).

**Fig.7.**
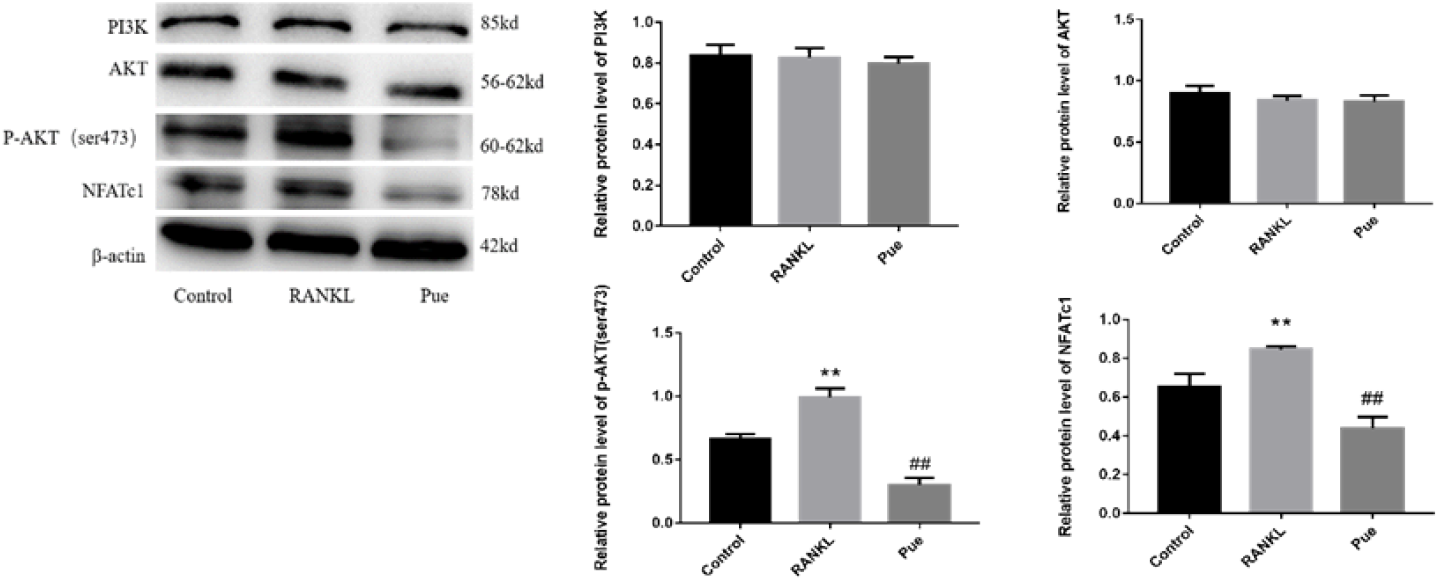
Western blotting measurement of Puerarin treatment on PI3K-AKT signaling pathway. Puerarin inhibited the activation of P-AKT (ser473) and down-regulated the expression of NFATc1. Data are expressed as the mean ± S.D. ** *P* < 0.01 vs control group, ## *P* < 0.01 vs induction group.

### Effects of puerarin on the protein expression of FoxO1, P-FoxO1, Catalase

We used Western Blot analysis to detect the effect of puerarin on FoxO1, P-FoxO1, Catalase protein expression. Compared with the RANKL group, the protein expression levels of FoxO1 and Catalase were significantly up-regulated and the protein expression levels of P-FoxO1 were down-regulated after puerarin intervention. The above studies suggest that puerarin may inhibit the proliferation and differentiation of osteoclasts by increasing the transcriptional activity of FoxO1 and increasing the expression of Catalase (Fig. 8).

**Fig. 8.**
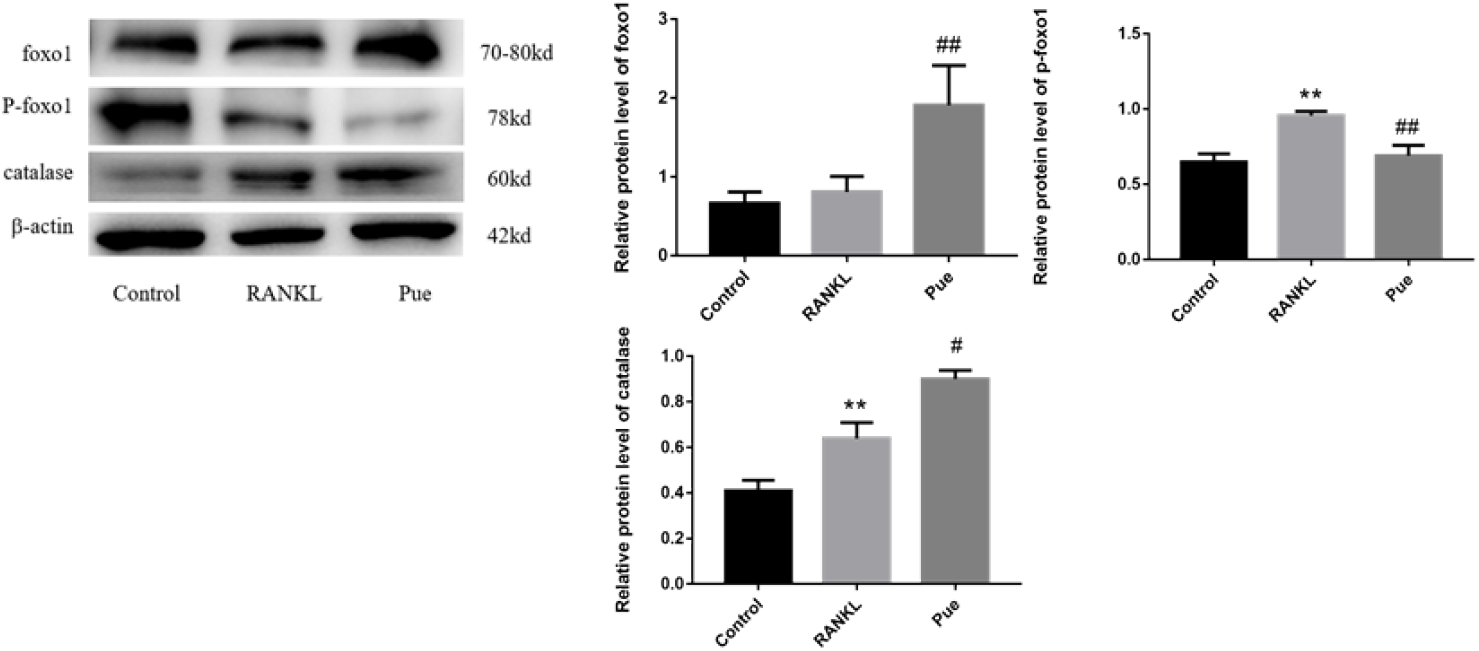
Western blotting measurement of Puerarin treatment on FoxO1, P-FoxO1 and catalase. Puerarin increased the expression of FoxO1 and catalase. Data are expressed as the mean ± S.D. ** *P* < 0.01 vs control group, # P < 0.05, ## P < 0.01 vs induction group.

## DISCUSSION

An increased ROS production or a decreased antioxidant activity is the main cause of oxidative stress in biological systems. The production and release of pro-inflammatory factors are increased when ROS activate related signaling pathways, perpetuating the inflammatory response. Nevertheless, excessive stimulation or prolonged inflammation in the body can damage tissues, causing many diseases such as periodontitis, bronchitis, hypertension, tumors, and osteoporosis[12]. A transcription factor called forkhead box O transcription factor 1 (FoxO1) regulates bone mass by up-regulating antioxidant enzymes such as catalase and superoxide dismutase (SOD). The primary functional classifications of FoxO target genes are “stress response and antioxidant defense,” “metabolism,” and “cell death and proliferation”[13], although the antioxidant impact is the most important. Phosphorylation, acetylation, and ubiquitination of distinct FoxO protein sites by various cytokines or signal pathways impact the transcriptional activity, stability, and subcellular localization of FoxO protein [14,15]. The most important regulation mode is the phosphorylation modification of AKT. Translocation of FoxO protein to the nucleus occurs as a result of stress or growth factor deficiency. Alternatively, when the PI3K-AKT signaling pathway is persistently activated in cancer cells, AKT phosphorylates the FoxO transcription factors, which translocate into the cytoplasm for inactivation[16]. From the standpoint of antioxidants, we investigated the relevance of the PI3K-AKT signaling pathway in the effects of puerarin on osteoclast proliferation, activity, and function. By modulating the PI3K-AKT signaling pathway, puerarin was discovered to enhance the transcriptional activity of Foxo1 and raise catalase protein expression level and exhibit its antioxidant function to limit the proliferation and differentiation of osteoclasts. Resveratrol might upregulate the transcriptional activity of FoxO1 by blocking the PI3K-AKT signaling pathway, according to Yanling Feng [17], boosting resilience to oxidative stress and decreasing osteoclastogenesis. This finding provides a theoretical and experimental basis for the clinical treatment of pathology from the attitude of antioxidants.

In order to hold out higher analysis, we have a tendency to establish an osteoclast model as the next research object. RAW264.7 can differentiate into osteoclast-like cells in the presence of RANKL and M-CSF. Compared with mouse bone marrow macrophages (BMM), RAW264.7 cells have the benefits of straightforward in vitro culture and passage, easy induction into osteoclasts, and no need for isolation and purification. Our findings further showed that after being induced and cultivated for 4-6 days in an induction medium containing 20 ng/mL M-CSF and 50 ng/mL RANKL, RAW264.7 cells could generate osteoclasts. We discovered that the condition of RAW264.7 cells influences the production of osteoclasts when growing them. The cell morphology becomes uneven, the antennae expand, and the differentiation is severe when the cells are subcultured too much. Even boosting the induction solution’s M-CSF and RANKL cytokines or lengthening the induction duration won’t stop RAW264.7 cells from effectively transforming into osteoclasts at this point. The induction period will be lengthened if RAW264.7 cells are solely given RANKL-containing media instead of M-CSF-containing medium. Even if RANKL-containing media is continuing to be supplied, too much M-CSF will cause the cells to proliferate excessively thickly eventually fail to differentiate into osteoclasts either. According to Chengchao Song[19], the appropriate cell seeding density and RANKL intervention time point affect the number of osteoclasts formed, which is consistent with our findings: when the cell density is too small or too large, neither can successfully induce RAW264.7 cells to become osteoclasts. Therefore, it’s vital to work out the seed plate density before the beginning of the experiment for the booming induction of osteoclasts. In the following study, it was found that puerarin might inhibit the differentiation of osteoclasts, cut back the generation of osteoclasts and inhibit the expression of osteoclast marker genes TRAP, MMP-9, and CTSK by TRAP staining, cell counting and qPCR experiments. In vitro studies demonstrated that Puerarin regulates ERK1/2-runx2 signaling pathway to influence the differentiation of BMSCs into osteoblasts, as well as inhibition of autophagy and proliferation of osteoclast precursors to inhibit osteoclast differentiation[10].

The PI3K-AKT signaling pathway is critical for cell growth, proliferation, and differentiation[20, 21]. When cells are stimulated by a signal, PIP3 produced by PI3K allows AKT with a PH domain to transfer to the cell membrane, where it is phosphorylated at the threonine phosphorylation site (Thr308) and serine phosphorylation site (Ser473) of the AKT protein by 3-phosphoinositide-dependent protein kinase 1 (PDK1) and 3-phosphoinositide-dependent protein kinase 2 (PDK2). Phosphorylation-activated AKT also impacts downstream target proteins, allowing it to carry out its roles of cell growth, proliferation, and apoptosis inhibition[22]. The phosphatase and tensin homolog gene (PTEN), which was lost from chromosome 10, possesses both lipid and protein phosphatase activity. Phosphatidylinositol 3 phosphate (PIP3) is dephosphorylated to phosphatidylinositol 2 phosphate (PIP2), which suppresses protein kinase B (PKB/AKT) phosphorylation activation (p-AKT) and so adversely regulates the PI3K-AKT signaling pathway[23]. During osteoclast development, PTEN plays a function in regulating the PI3K-AKT signaling pathway [24-26]. Catalpol, a bioactive molecule derived from Rehmannia glutinosa that has anti-inflammatory, antioxidant, anti-diabetic, and anti-tumor properties, blocked RANKL-induced activation of the NFB and AKT signaling pathways, as well as downregulating the expression of NFATc1 and downstream proteins. Catalpol blocking the ubiquitination and degradation of PTEN and boosting the protein expression level of PTEN may be the mechanism preventing osteoclast differentiation[27]. Combining our findings with earlier research, we discovered that puerarin can inhibit the PI3K-AKT signaling pathway by raising PTEN expression and promoting catalase production, therefore alleviating and curing osteoporosis. As a result, we wondered if puerarin may alter the PI3K-AKT signaling pathway by modulating PTEN activity during RANKL-induced osteoclastogenesis. Unfortunately, we discovered no significant difference in PTEN before and after puerarin intervention in our investigation, indicating that PTEN may not play a significant role in blocking the activation of the PI3K-AKT signaling pathway during osteoclast generation. PTEN is controlled by a range of alterations, including phosphorylation, acetylation, and ubiquitination, which should all be evaluated [28-31]. In addition to osteoclasts, bone tissue contains osteoblasts, osteocytes, osteoprogenitors, and other cells. The next stage will be to investigate the function of PTEN in different cells.

## Acknowledgments

This work was supported by National Natural Science Foundation of China(81560362 and 82160550)

## Compliance with enthical standards

Conflict of Interest

The authors state that there is no conflict of interest regarding this manuscript. Ethical statement

There are no animal expriments carried out for this article.

## Contributor information

Jisheng Xie, Yiqiu Yang

